# Understanding the genetic risks of complex diseases using the additive epistatic interaction model: a simulation study

**DOI:** 10.1101/865196

**Authors:** Yupeng Wang

**Author notes:** Corresponding author: Yupeng Wang, Ph.D.

## Abstract

Thousands of Genome-Wide Association Studies (GWAS) have been carried out to pinpoint genetic variants associated with complex diseases. However, the proportion of phenotypic variance which can be explained by the identified genetic variants is relatively low, leading to the “missing heritability” problem. This problem may be partly caused by the inadequate understanding of the genetic mechanisms of complex diseases. Here, we propose the additive epistatic interaction model, consisting of widespread pure epistatic interactions whose effects are additive and can be summarized by a genetic risk score. Based on a simulated genotype dataset, the additive epistatic interaction model well depicted genetic risks and hereditary patterns of complex diseases. Based on the 1000 Genomes Project data, the additive epistatic interaction model accurately classified human populations. Moreover, the model’s genetic risk score can be replaced by a deep learning model which is more resistant to noises. We suggest that the additive epistatic interaction model may help to understand the genetic mechanisms and risks of complex diseases.

## Introduction

Thousands of genome-wide association studies (GWAS) have been carried out to identify genetic variants associated with complex diseases or quantitative traits [1-3]. In a GWAS, several to hundreds of genetic variants may be reported. However, they can explain only a small proportion of heritability, which is often referred to as the “missing heritability” problem [4]. Various theories have been proposed to explain the “missing heritability” problem [5]. One notable theory is epistatic interactions, referring to the phenomenon that genetic variants may show little or moderate individual effects, but strong interaction effects [6, 7]. Other theories such as rare variants [8, 9] and gene-environment interactions [10] have been also widely investigated.

Complex diseases and quantitative traits are often determined by multiple genetic variants [11, 12]. Genetic risk score (GRS), or polygenic score, which adds up the impacts of multiple genetic variants, has become a popular post-GWAS analysis approach [13, 14]. Utilization of GRS has led to summarized disease risks leveraging multiple genetic variants and environmental factors [15-17], which often show higher predictive power than using individual genetic variants. However, previous applications of GRS mainly included genetic variants with marginal effects. Therefore, GRS for epistatic interactions is underexplored. Although aggregated scores of Multifactor Dimensionality Reduction (MDR) were investigated years ago [18, 19], the computational burden of MDR has rendered it impossible to perform genome-wide scan of epistatic interactions. Because genome-wide genotype datasets, especially those produced by next-generation sequencing technologies, are being increasingly generated, efficient models for summarizing the genetic risks of epistatic interactions are highly desirable.

To date, most computational tools for detecting epistatic interactions actually capture those epistatic interactions with marginal effects. However, in pure epistatic interactions, individual genetic variants have zero marginal effect [20, 21]. Due to computational difficulties, genetic variants involved in pure epistatic interactions were seldom reported in previous GWAS. However, pure epistatic interactions could be an important genetic component of common traits and diseases, which may substantially account for the “missing heritability”. Here, we infer that pure epistatic interactions are widespread in the human genome, and the effects of multiple pure epistatic interactions are in general linear and additive. We refer to this genetic model as the additive epistatic interaction model. Under this model, any individual has a genetic risk score which is the number of occurrences of interacting genotypes. We use simulated genotype datasets and 1000 Genomes Project data to show that the additive genetic interaction model can be useful and insightful for understanding the genetic mechanisms and risks of complex diseases.

## Methods

### Data simulation

We simulated a genotype dataset consisting of 500 pairs of interacting SNP loci containing pure epistatic interactions, and 1000 case and 1000 control individuals. For computational simplicity, we placed interacting loci at adjacent positions. All loci were assumed to be independent. The algorithm is described as follows:

1. *M* denotes the number of pairs of interacting loci, and *N* denotes the number of individuals in either the case or control group. The total number of loci is 2\**M* and total number of individuals is 2\**N*.
2. According to a uniform distribution between 0.01 and 0.5, assign a guiding MAF to each of the 2\**N* loci. The guiding MAFs are the same between case and control groups. This ensures that no marginal effect will exist at each of the 2\**N* loci.
3. Genotypes of any locus (major allele **A** and minor allele **a**) are generated according to its guiding MAF. For each allele of a genotype, a random number between 0 and 1 is generated according to a uniform distribution. If the random number is smaller than the guiding MAF, the minor allele **a** is generated. This procedure is repeated for all individuals in either the case or control group.
4. For the control group in which there is no interaction effect, the genotypes of 2\**M* loci are just generated according to step 3).
5. For the case group, each pair of adjacent loci composes interacting loci, and the occurrence of interaction effects is determined by the **aabb** genotype. For each pair of interacting loci, a large number (e.g. 5000) of genotype sets are generated, and the genotype set with the highest occurrences of **aabb** genotype is chosen as the final genotypes.

Based on the above simulated genotype dataset, the genotypes of three offspring groups were generated. For each group, 1000 individuals were generated. Random mating and zero mutation rate were assumed.

To add noises to the simulated genotype dataset, additional loci were generated according to the procedure for the control group. In the noise data, adjacent loci were assumed to be interacting and involved in computing GRS.

### Computing GRS

For each individual, the GRS is the count of **aabb** genotype among all pairs of interacting loci.

### Analysis of 1000 Genomes data

Genotypes and alternative allele frequencies (AAFs) of chromosome 20 from the 1000 Genomes Project were downloaded from the NCBI ftp site (ftp://ftp-trace.ncbi.nih.gov/1000genomes/ftp/release/20130502/). We built two genotype datasets containing pure epistatic interactions for each pair of super populations, by exchanging case and control assignments between the two super populations. Thus, a total of 20 genotype datasets were built. For each pair of super populations, we first selected the SNPs with 0.01≤AAF<0.5 in both super populations, which ensured that the alternative allele was always the minor allele at each locus. Next, we filtered out the SNPs with different MAFs between the two super populations (*p*≤0.05) according to χ^2^ test, which ensured that the MAF at each locus was similar between the two super populations. The remaining SNPs were the pool for generating 500 pairs of loci containing pure epistatic interactions. The algorithm of generating the 500 pairs of loci is described as follows:

1. Randomly select two loci which are separated by at least 10 loci, which ensures that putative interacting loci are not under high linkage disequilibrium.
2. Examine whether the selected two loci have more occurrences of **aabb** genotype code in the case group than the control group, according to χ^2^ test.
3. If *p*<0.01, the selected two loci compose a pair of interacting loci, and are added to the genotype dataset containing pure epistatic interactions and excluded from the SNP pool; otherwise, the two loci are put back to the pool.
4. The above procedure is repeated until all 500 pairs of interacting loci are filled.

### Deep learning models

Deep learning was performed using Tensorflow and Keras, available via Python 3.5.2. Deep neural network (DNN) models were built to classify case and control statuses. The structure of the DNN contains:1) layer 1: dense, 512 units, relu activation, 0.25 dropout; 2) layer 2: dense, 512 units, relu activation, 0.25 dropout; and 3) layer 3 (outcome): dense, 2 units, softmax activation. DNN models were trained based on the following parameters: optimizer: Adam; loss: categorical cross entropy; learning rate: 0.0005; training epochs: 20; and batch size: 256. The Sklearn python library was used to compute AUCs. Each analyzed dataset was divided into 80% training and 20% testing data. Five-fold cross validation was employed to generate an average AUC for each analyzed dataset.

### Availability

Source code is publicly available at https://github.com/wyp1125/additive_epistasis.

## Results

Simulation is an effective approach for evaluating new genetic models [22]. We simulated a case-control genotype dataset consisting of 1000 SNP loci and 2000 individuals (see Methods). The 2000 individuals were divided into case (i.e. disease) and control groups of equal size. Genotypes were generated according to the same guiding minor allele frequencies (MAFs) between the case and control groups. In the case group, the 1000 loci composed 500 pairs of interacting loci containing pure epistatic interactions. Therefore, all the 1000 loci should be regarded as association with the disease. The alleles of a pair of interacting loci are denoted by A/a and B/b (**a** and **b** are minor alleles) respectively. We assume that only the **aabb** genotype (i.e. both loci are homozygous for their minor alleles) has interacting effects. Under conventional genetic models, the case group has higher disease allele frequencies than the case group, whereas in this simulated dataset, no locus displays significantly different allele frequencies between the case and control groups, because the minimum *p-*value (χ^2^ test) is 7.03×10^−5^ which is higher than the standard GWAS cutoff of 5×10^−8^ as well as the lenient cut-off of 10^−5^ [23] and no locus leads to a significant *p*-value after FDR adjustment. The mean and median relative risks of the **aabb** genotype among all pairs of interacting loci are 1.63 (>1, *p*<2.2×10^−16^, *t*-test) and 1.60 respectively, confirming that pure epistatic interactions were successfully generated.

The genetic risk score (GRS) of an individual is defined as the number of occurrences of the **aabb** genotype among the 500 pairs of interacting loci. We computed the GRS for each case and control individual. Distributions of GRS (Figure 1) clearly show that the case group tends to have higher GRS than the control group (*p*<2.2×10^−16^, *t*-test). Although the above comparison is informative, clinical values of GRS depend more on whether the GRS can distinguish case (i.e. disease) and control (i.e. healthy) statuses. To this end, we computed the area under the ROC curve (AUC), a measure of the discriminative ability ranging from 0 to 1. An AUC of 0.900 was obtained, indicating that the GRS can well depict disease risks.

**Figure 1.**
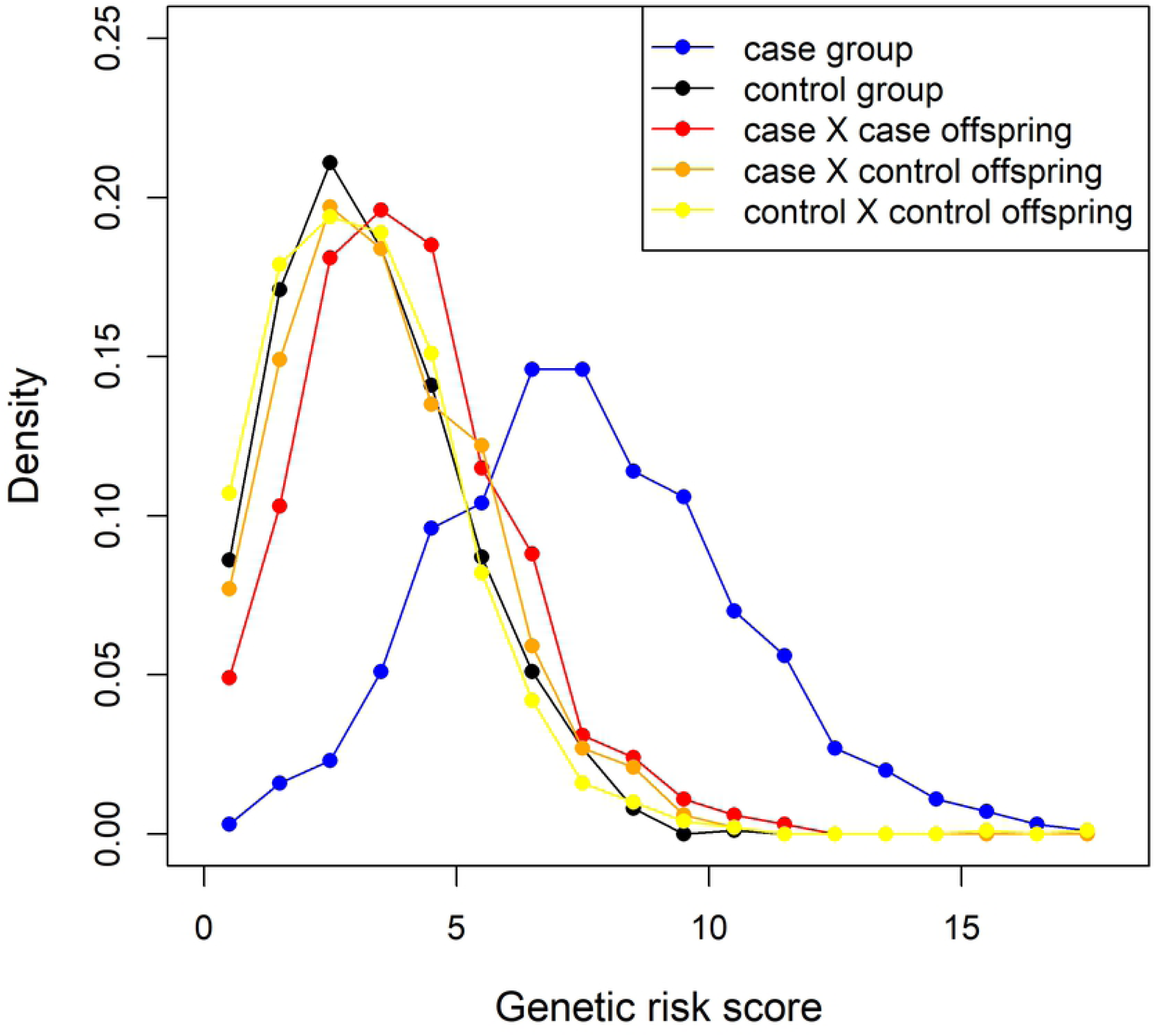
Comparisons of GRS distributions among case, control and different offspring groups.

The exact hereditary mechanisms of complex diseases such as cancer and heart diseases are still little understood. Population and clinical studies suggest that the offspring of healthy parents tend to have less risks of developing a complex disease than the offspring of one parent or both parents with histories of that complex disease [24-26]. We asked whether offspring’s risks of developing a complex disease can be depicted by the GRS under the additive epistatic interaction model. Based on the simulated genotype dataset, we derived the genotypes of offspring belonging to three parent groups: 1) case×case, 2) case×control, and 3) control×control (see Methods). We added the GRS distributions of the three offspring groups to Figure 1. The case × case offspring tend to have higher GRS than the other two offspring groups (*p*=2.47×10^−7^ and *p*<2.2×10^−16^ respectively, *t*-test), and the case × control offspring tend to have higher GRS than the control × control offspring (*p*=1.21×10^−4^, *t*-test). This analysis suggests that stronger parental histories of a complex disease do increase offspring’s genetic risks of developing that disease. However, the GRS of case × case offspring are greatly reduced relative to the case group (*p*<2.2×10^−16^, *t*-test), suggesting that even with strong parental histories of a complex disease, the genetic risks of developing that disease in offspring are much lower than the same genomes as real patients, because recombination significantly reduces the occurrences of the **aabb** genotype. This observation further indicates that for people with strong family histories of a complex disease, the actual acquirement of that disease may still largely depend on non-parental factors such as accumulation of de novo/somatic mutations and epigenetic modifications. The above analyses collectively suggest that the additive epistatic interaction model and its GRS can well depict the genetic risks and hereditary patterns of complex diseases.

Next, we asked whether the additive epistatic interaction model is applicable to real genotype data. To this end, we obtained the genotype data of chromosome 20 from the 1000 Genomes Project [27]. The genotype data consisted of 2504 individuals from 5 super populations including African (AFR), Ad Mixed American (AMR), East Asian (EAS), European (EUR) and South Asian (SAS). Here, we treat these super populations as the phenotype, and any two super populations comprise a pair of case-control groups. We inferred that if pure epistatic interactions are widespread in the human genome, the additive epistatic interaction model can be applied to classify super populations. We employed a customized procedure (see Methods) to identify 500 pairs of SNP loci containing pure epistatic interactions for each pair of super populations. Next, we computed AUCs for pairs of super populations based on the GRS of 500 pairs of interacting loci. Note that for each pair of super populations, we exchanged case and control assignments, resulting in two AUC values. These AUCs range from 0.866 to 0.991 (Figure2), suggesting that human super populations can be accurately classified based on the additive epistatic interaction model.

**Figure 2.**
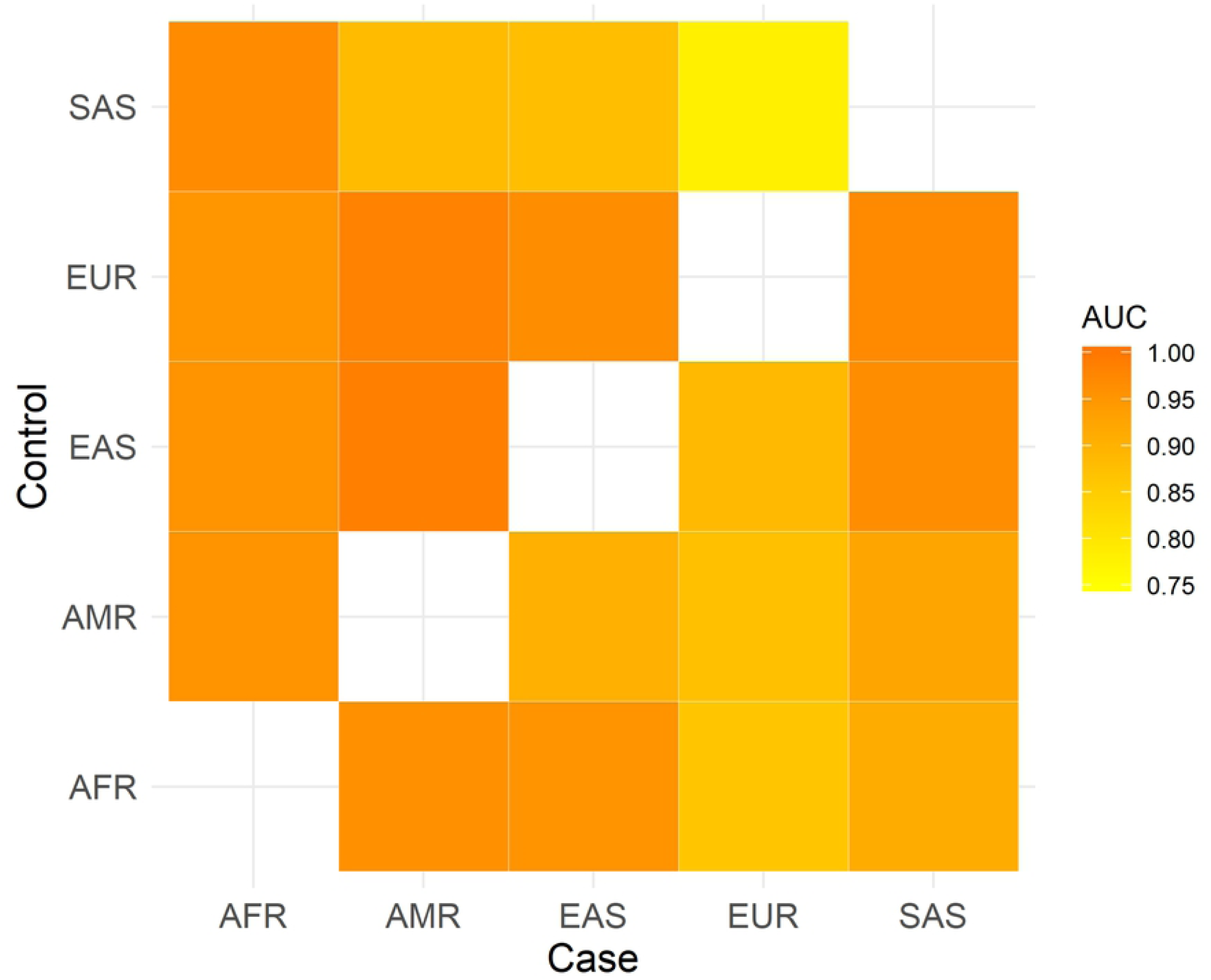
AUCs of using the additive epistatic interaction model to classify human super populations.

An important consideration for applying the additive epistatic interaction model is its robustness. In real data applications, especially those utilizing whole-genome sequencing technologies, the numbers of markers are much larger than sample sizes. Thus, a part of the identified pairs of interacting loci may not be real. Therefore, we asked whether the additive epistatic interaction model can tolerate some extent of noises and whether advanced machine learning technologies such as deep learning can be integrated to the additive epistatic interaction model. While deep learning has different models, we chose deep neural network (DNN) because different pure epistatic interactions do not contain sequence or interdependency information. We here note that the features of DNN models should be occurrences of the **aabb** genotype, rather than original genotypes because effective feature extraction is still required for deep learning technologies.

We added different ratios of noises (i.e. loci assumed to be interacting but do not contain pure epistatic interactions) to the simulated genotype dataset, and computed AUCs of distinguishing between case and control statuses based on GRS and DNN respectively. Figure 3 shows comparisons of AUCs between GRS and DNN at different noise ratios. In general, the AUCs of both approaches gradually decrease along with increase of noise ratios. When noise ratios are relatively low (e.g. <1), GRS outperforms DNN. When the noise ratio goes up from 1, the AUCs of GRS decrease rapidly, while the AUCs of DNN show resistance to noises, leading to DNN outperforming GRS. This analysis suggests that both GRS and DNN can tolerate a substantial proportion of noises, and DNN can be integrated to the additive epistatic interaction model for dealing with complex datasets.

**Figure 3.**
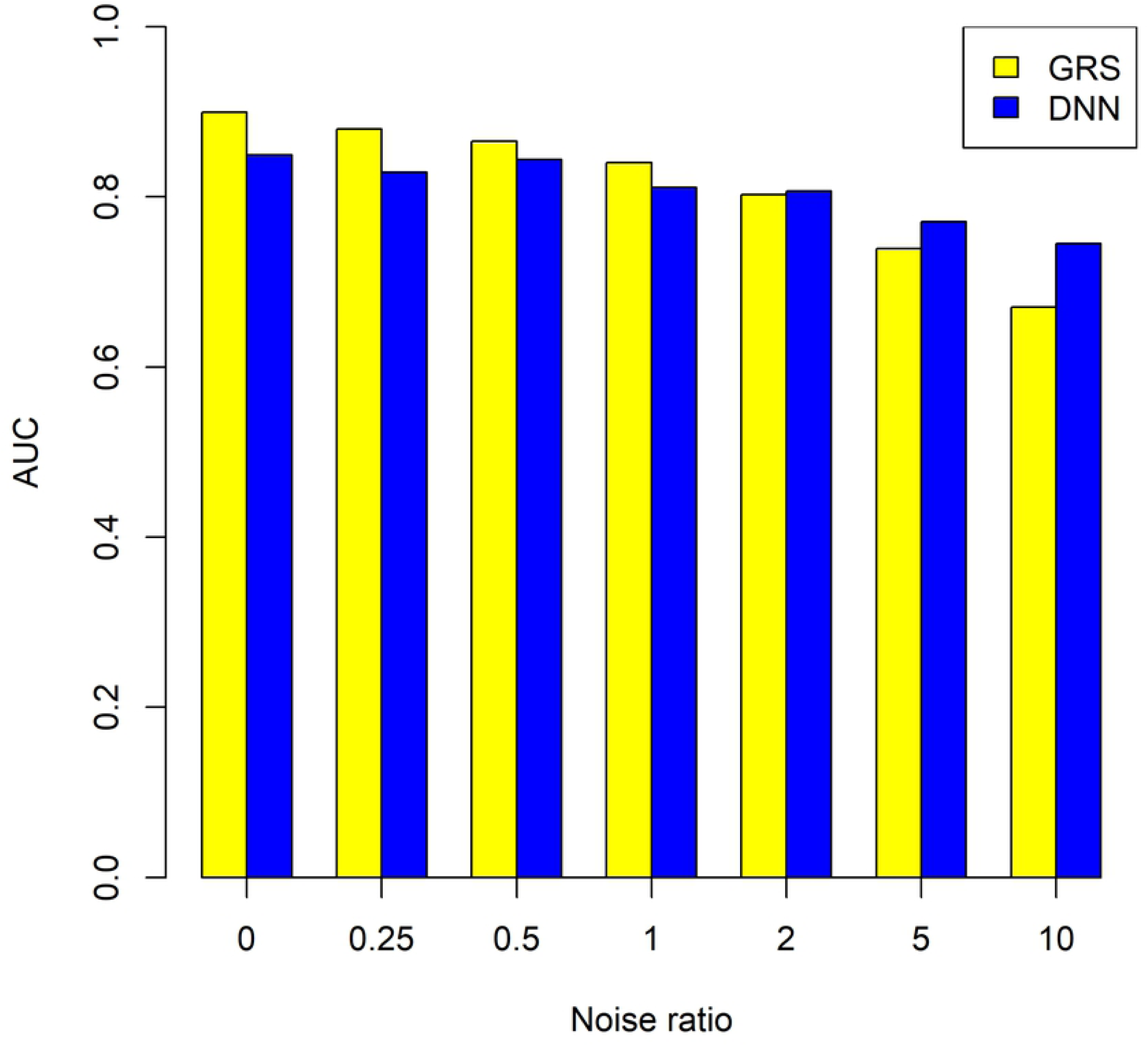
Comparisons of AUCs between GRS and DNN at different noise ratios.

## Discussion

In this study, we propose the additive epistatic interaction model. We show that the model well depicts the genetic risks and hereditary patterns of complex diseases, and it is effective in classifying human populations. We also show that deep learning can be integrated to the additive epistatic interaction model for dealing with complex genotype datasets. Thus, the additive epistatic interaction model can be useful for understanding the genetic mechanisms and risks of complex diseases. In addition, we suggest that the extent of recombination is the key factor for reducing the GRS of the additive epistatic interaction model.

The underlying genetics of complex diseases are very complicated and a single genetic model may not capture the whole picture. Individual variants with strong marginal effects were most frequently investigated and reported in previous GWASs. Rare variants, which can be reliably assayed using exon sequencing technologies, are increasingly studied in recent years [28, 29]. In contrast, pure epistatic interactions have been seriously underexplored. In this study, we show that pure epistatic interactions can be widespread in the human genome, and human populations can be accurately classified based on the additive epistatic interaction model. This result indicates that pure epistatic interactions should be investigated in future genetic association studies, and re-analysis of previous GWAS using the additive epistatic interaction model may be also insightful.

To classify human super populations, we adopted a random selection algorithm which selected only 500 pairs of loci containing pure epistatic interactions from chromosome 20. This selection procedure took about 15 minutes based on a small Linux workstation with a CPU of 3.10GHz×4 and memory of 16G. The number of genetic variants in chromosome 20 was∼1.8M, which is higher than typical numbers of loci in GWAS, suggesting that pure epistatic interactions can be rapidly assessed in GWAS, and an exhaustive search may be feasible considering that high-performance super computers are widely available in research institutions. We believe that the AUCs of the additive epistatic interaction model can be further increased if interacting loci are selected by exhaustive search and top statistical significance.

Assessing pure epistatic interactions in this study has involved rare variants with 0.01<MAF<0.05. If the sample size is big enough (say >10k), rare variants with even smaller MAFs can be included. Complex diseases may be affected by both common variants and rare variants/mutations. The “missing heritability” should not be a real problem considering that there are still many limitations on both experimental and data analysis sides. Investigation of the interplays among different genetic models/components is highly desirable in future genetic association studies for complex diseases and quantitative traits.

## Acknowledgements

I thank Ying Sun and Xinyue Liu for valuable discussion of this research.

